# CIDER: a pipeline for detecting waves of coordinated transcriptional regulation in gene expression time-course data

**DOI:** 10.1101/012518

**Authors:** Marco Mina, Giuseppe Jurman, Cesare Furlanello

## Abstract

Cell adaptability to environmental changes is conferred by complex transcriptional regulatory networks, which respond to external stimuli by modulating the expression dynamics of each gene. Hence, deciphering the network of transcriptional regulation is remarkably important, but proves to be extremely challenging, mainly due to the unfavorable ratio between the number of available observations and the number of parameters to estimate. Most of the existing computational methods for the inference of transcriptional networks consider steady-state gene expression datasets, and produce models of transcriptional regulation best explaining the observed static gene expression.

Gene expression time-courses are an emergent typology of gene expression data, paving the way to the characterization of the time-dependent dynamics of transcriptional regulation.

In this work we introduce the Complexity Invariant Dynamic Time Warping motif EnRichment (CIDER) analysis, a novel computational pipeline to identify the prominent waves of coordinated gene transcription induced in cells by external stimuli, and determine which TFs are involved in the coordination of gene transcription. The CIDER pipeline combines unsupervised time series clustering and motif enrichment analysis to first detect transcriptional expression patterns, and then identify the TFs over-represented in the promoter regions of gene sets with similar expression dynamics.

The ability of CIDER to correctly identify regulatory interactions is assessed on a realistic synthetic dataset of gene expression timecourses, generated by simulating the effects of knock-out perturbations on the E. coli regulatory network.

The CIDER source code and the validation datasets are available on request from the corresponding author.

## 1 INTRODUCTION

Cells respond to external stimuli and adapt to environmental changes by modulating their gene expression through complex regulatory programs. Not surprisingly, the disregulation of these regulatory programs plays a major role in complex diseases, such as cancer and autoimmune diseases. Transcriptor Factors (TFs), molecules with the ability of selectively bind to specific DNA regions and enhance/inhibit the transcription of neaby genes, are among the most studied regulators of gene expression. Consequently, current approaches to improve cancer treatment try to identify the proper combination of TFs that would induce the desidered gene expression when opportunely blocked or induced.

TFs do not operate separately, as their transcriptional effects combine in complex patterns to orchestrate and synchronize the transcription of genes. The ensemble of the regulatory interactions taking place within cells is commonly referred to as transcriptional regulatory network (TRN). Deciphering TRNs is an essential step to characterize the role of each TF, its involvement in specific diseases, and the potential therapeutic effects of targeting it. The task is challenging due to the small amount of available data, compared to the number of parameters to estimate and the noise introduced by the experimental procedures. Morevoer, TF activity within the regulatory networks is cell- and time-specific, and the TRNs can vary significantly even within the same cell under different external stimuli.

The typical approach to understand TRNs consists of collecting the steady-state gene expression of cells under a specific condition (e.g. stimulated with a particular factor) and build a model of transcriptional regulation that best explain the observed gene expression. The inferred regulatory mechanism is a static description of the regulatory effects of the TFs on the target genes, critical for understanding the condition-specific role of TFs. Concretely, the typical output of the proposed methodologies is a network of regulatory interactions connecting TFs to the regulated genes [1], [2]. Additional data such as the strength, direction, and type of regulation are provided by some of the methods.

Transcriptomics time-courses (e.g. microarray or RNASeq) are a longitudinal representation of gene expression useful for characterizing the dynamics of regulatory activity within cells. Transcriptomics time-courses describe the evolution of gene expression within the profiled cells across time-points, thus extending the static description provided by steady-state experiments [3], [4]. Hence, analyzing time-course data can help characterizing the mechanisms of transcriptional regulation triggered by external stimuli. By analyzing the time-courses of gene transcription initiation, for instance, Arner et al. showed that external stimuli propagate along the transcriptional regulatory networks by inducing waves of coordinated transcription characterized by specific expression patterns [5]. Current network inference methodologies are partially fit to characterize these transcriptional waves, as they generally try to infer the single regulatory interactions, rather than the coordinated regulation of multiple genes.

In this work we introduce the Complexity-Invariant Dynamic Time-Warping motif EnRichment (CIDER) analysis, a computational pipeline to detect the prominent waves of coordinated transcription in time-course data, and infer which TFs are involved in the underlying regulatory mechanisms. CIDER differs from current approaches as it focuses on identifying the regulatory effects of TFs on groups of genes with similar expression dynamics, rather than trying to dissect the single regulatory interactions.

The following sections describe CIDER methodology and implementation, and present an assessment of its performance on a realistic synthetic dataset.

## 2 CIDER

Conceptually, a wave of coordinated transcription consists of a set of genes whose expression patterns, under the regulation of one or more TFs, follow a specific dynamic in time. Reversing this statement, we reasoned that **if a group of genes with similar trajectories is regulated by a common set of TFs, then it qualifies as a transcriptional wave**. Following this idea, CIDER first detects clusters of genes with similar expression patterns from time-course data, and then determines whether there are TFs that consistently participate to the regulation of the genes clustered together. The clusters of genes with similar expression that are supported by evidence of consistent co-regulation are reported by CIDER as transcriptional waves.

Both the tasks of detecting clusters of genes with similar expression and of finding TFs significantly regulating the clustered genes present a series of difficulties that must be tailored with the utmost attention. First of all, the increment of data-points on the temporal scale is generally counter-balanced by a significant drop in the number of independent samples available. Moreover, the longitudinal nature of the time-course data introduce an inter-dependency between different time-points. Hence, the methodologies for gene clustering must be adapted.

CIDER is designed to require just a single time-course from a single biological replicate. Provided that the number and arrangement of time-points is sufficient to discern the expression patterns, CIDER opens the possibility of analyzing transcritpional regulation at singe-sample resolution. Obviously, time-courses from multiple biological replicates can be joined to improve the clustering quality and stability.

On the other side, inferring the regulatory interactions between TFs and regulated genes is a long-standing problem for which different solutions, more or less effective, have been proposed. While a group of solutions rely only on the expression data, others integrate other information to improve the accuracy of the inference. Both families of solutions have been successfully used to infer regulatory network. We propose to consider external data to improve the inference accuracy in some cases, such as when no correlation between TF and target expression is present. This happens, for instance, when a TF is constitutively expressed but inactive, and begins its regulatory activity after a post-translational modification. CIDER makes use of a-priori known information about TF binding motifs to evaluate the transcriptional effect of each TF on each single cluster.

### 2.1 Algorithm description

CIDER consists of two steps: first, expression patterns are identified by unsupervised hierarchical clustering of the time series. Then, motif enrichment analysis is performed on the time series clusters, to identify the transcription factors likely to induce the transcription of the genes with similar expression pattern. The pipeline, sketched in Fig. 1, is described in the rest of this section from both the methodological and implementative points of view.

- **Required input data**

Three different types of data are required by CIDER:

1. The time series describing gene expression longitudinally across time, used to cluster genes with similar longitudinal expression patterns,
2. A collection of TF binding sites, whose over-representation in the promoter regions of coclusterer genes (motif enrichment analysis) is interpreted as indicator of TF transcriptional activity,
3. The reference genome, required to extract the flanking sequences of co-clustered genes and the background sequences required by the motif enrichment analysis.

- **Step 1. Time series clustering**

A hierarchical clustering approach is adopted to identify expression patterns from the longitudinal gene expression data. The following paragraphs provide details on preprocessing, distance, and specific hierarchical clustering methods adopted in CIDER.

*Step 1-A. Time series preprocessing*: time series are constrained between 0 and 1 by dividing each time series by its maximum value, in order to apply distances that require time series normalization (see step 1-B). Each replicate is normalized separately, to avoid penalizing replicates with systematic lower signal levels (e.g. low tag per million counts), and focus on shape patterns instead of magnitude levels. To reduce the impact of outliers in short time series, instead of averaging the replicates, they are concatenated obtaining longer time series.

**Fig. 1:**
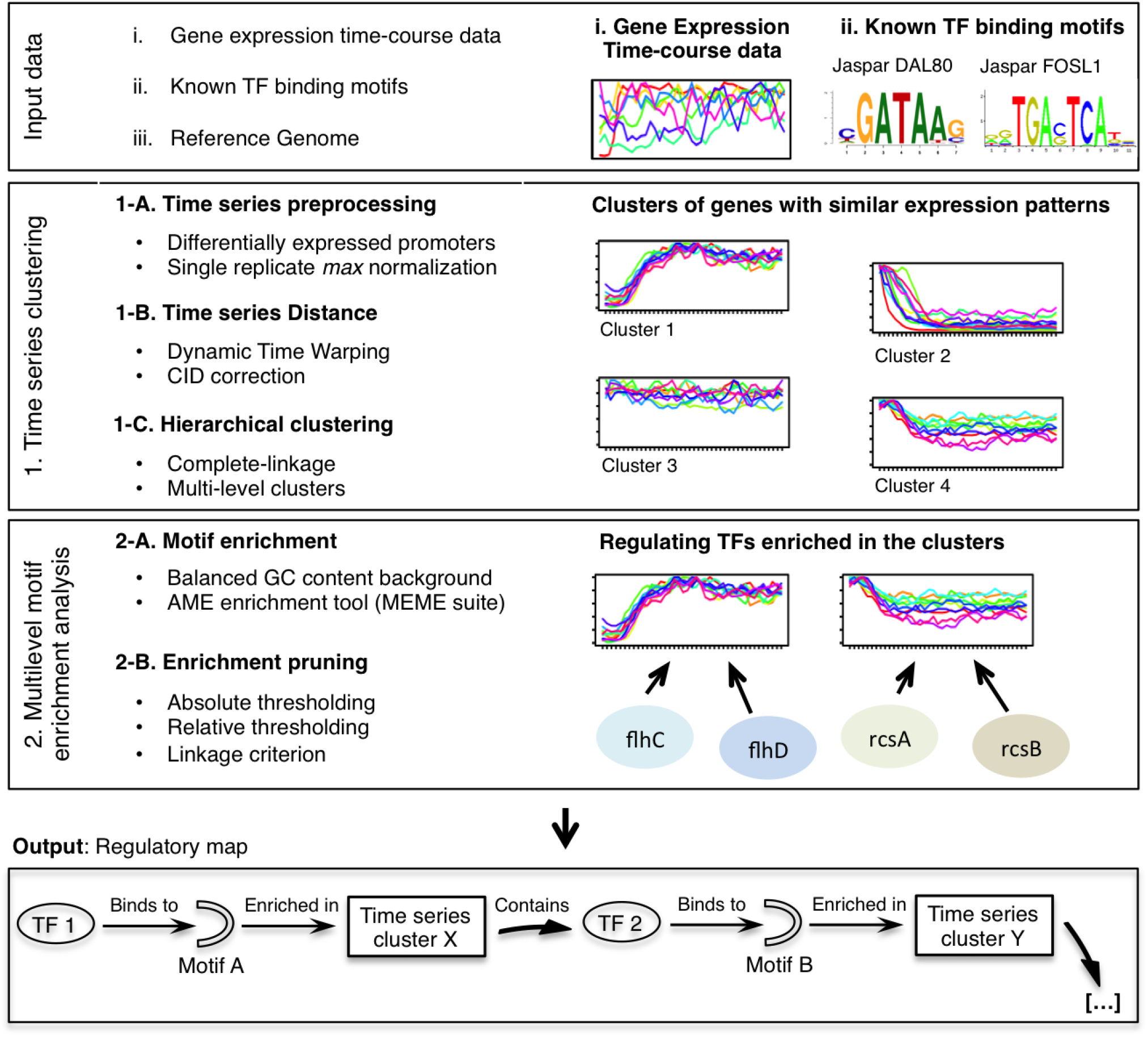
Overview of the CIDER pipeline. Time series clustering and multilevel motif enrichment analysis are combined to infer the regulatory map from time-course data and known TF binding motifs. The four TFs listed at step 2 (*flhC/D*, *rcsA/B*) are from the *E.coli* regulatory network (see Fig. 2).

*Step 1-B. Time series distance*: to overcome known issues with correlation-based methods [6], [7], as an appropriate distance for time series clustering, CIDER adopts the Euclidean-based Dynamic Time Warping (DTW) distance [8], The application of DTW on transcriptomics time series profiling has been already demonstrated on microarray platforms [9]. DTW alignment is constrained to align time-points within each replicate, with a Sakoe-Chiba band of width 1 as global constraint [10]. As a further refinement, we implemented in CIDER the Complexity Invariant (CID) correction for DTW (CIDDTW method), one of the latest installations of DTW, proposed as a generalization that accounts for signal complexity [11]. In details, the CIDDTW method introduces a correction term penalizing signals with low variability; the rationale is that for standard distance measures (Euclidean, correlation, non-corrected DTW) pairs of complex objects tend to be gauged further apart than pairs of simple objects, introducing errors in classification.

*Step 1-C. Hierarchical clustering*: Time series are clustered in CIDER by means of the complete-linkage agglomerative hierarchical clustering algorithm.

- **Step 2. Multilevel Motif Enrichment analysis**

CIDER adopts motif enrichment as the pruning rule for clustering. In general, motif enrichment is defined as a class of techniques to identify motifs with a significant number of binding sites in a set of DNA sequences (i.e. in the flanking regions of the transcription start sites of a group of genes).

*Step 2-A. Motif Enrichment*: The hierarchical clustering structure produced at step 1.C is used as starting point to perform a multi-level motif enrichment analysis. Each node in the cluster is tested for enriched motifs using the AME tool, part of the MEME software suite [12] with default parameters.

The flanking regions of each gene in the cluster are extracted from the Human Genome using SAMtools [13]. By default CIDER considers the -1500 to +500 bases as flanking region. Overlapping flanking regions are then merged in a single sequence to avoid overcounting binding sites. For each cluster, a 10-fold set of random sequences selected from the pool of flanking regions of all the genes in the genome is used as background set for motif enrichment. The GC content of query and background sets is matched to avoid the GC content bias in motif enrichment, as described in [14]. For each motif, AME counts the number of binding sites in the foreground and background sets of flanking regions, and evaluates the over-representation in the foreground set by Fisher’s exact test.

*Step 2-B. Enrichment-based pruning*: The enrichment associations computed by AME are then pruned to retain only the most relevant associations between motif activity and expression patterns. In literature, different approaches were proposed to prune not significant association [15], [16], [17]. CIDER includes three pruning strategies:

- Absolute thresholding: the association between a cluster and a motif is deemed as significant if the Fisher test P value is smaller than a given threshold *P*_*max*_ [15], [16].
- Relative thresholding: the association between a cluster and a motif is deemed as significant if the Fisher test P value is greater than *α* standard deviations the mean of the enrichment P values of the other transcriptional motifs in the cluster [17].
- Linkage criterion: and extension of the absolute thresholding requiring the enrichment to be redundant at multiple levels of the clustering hierarchy. This approach, inspired by the shadow analysis [18], further refines the enrichment by removing the motif-cluster associations that are not supported by contiguous enrichments along the clustering hierarchy. The rationale is that a cluster might be enriched for a specific motif even when only a subset of its genes is effectively regulated by the TFs binding to the enriched motif, while the other sub-cluster is not significantly enriched. Hence, the linkage criterion reduces false positives by ruling out sub-clusters that are not significantly enriched by themselves, but appears to be enriched in upper layers of the clustering hierarchy because merged with another significant cluster.

The three pruning strategies can be combined together to define stricter criteria.

### 2.2 Implementation

The time series clustering routines are entirely implemented as R scripts. The R package dtw [19] was extended to implement the CIDDTW, and the R package parallel used to parallelize the task on multiple processors. The R package fastcluster [20] is used to perform hierarchical clustering on thousands of time series, instead of the default R hclust hierarchical clustering solution.

The multi-level enrichment analysis module, instead, is series of Python, R, and bash scripts. In particular, a collection of bash scripts controls the AME enrichment procedure by running the AME application in parallel by using the GNU parallel utility [21].

The Python package ETE2 [22] was used to integrate clustering and enrichment data in an unified structure feasible for pruning.

### 2.3 Output description

The primary output provided by CIDER is a list of associations between time series clusters and TFs, as the one in Table 1. Extending the TF-cluster associations to the genes representated by each time series, it is possible to derive a broad regulatory map specific for the time-course. As a by-product, CIDER also provides the hierarchical clustering of the time series as Newick file.

## 3 VALIDATION

The validation approach commonly used to assess regulatory activity inference methods consists of comparing the inferred regulatory interactions and active TFs to the set of regulatory mechanisms known to take place in the tested dataset. Following this approach, we set up a procedure based on the idea of perturbing a (set of) TFs, collect the time series describing the resulting gene expression dynamics, and verify whether the perturbed TFs are included in those inferred by CIDER as involved in the regulation of a transcriptional wave. Finally, we also manually verified the soundness of the inferred transcriptional waves on a random selection of tests. The dataset generation and validation procedures are sketched in Fig. 2.

**Fig. 2:**
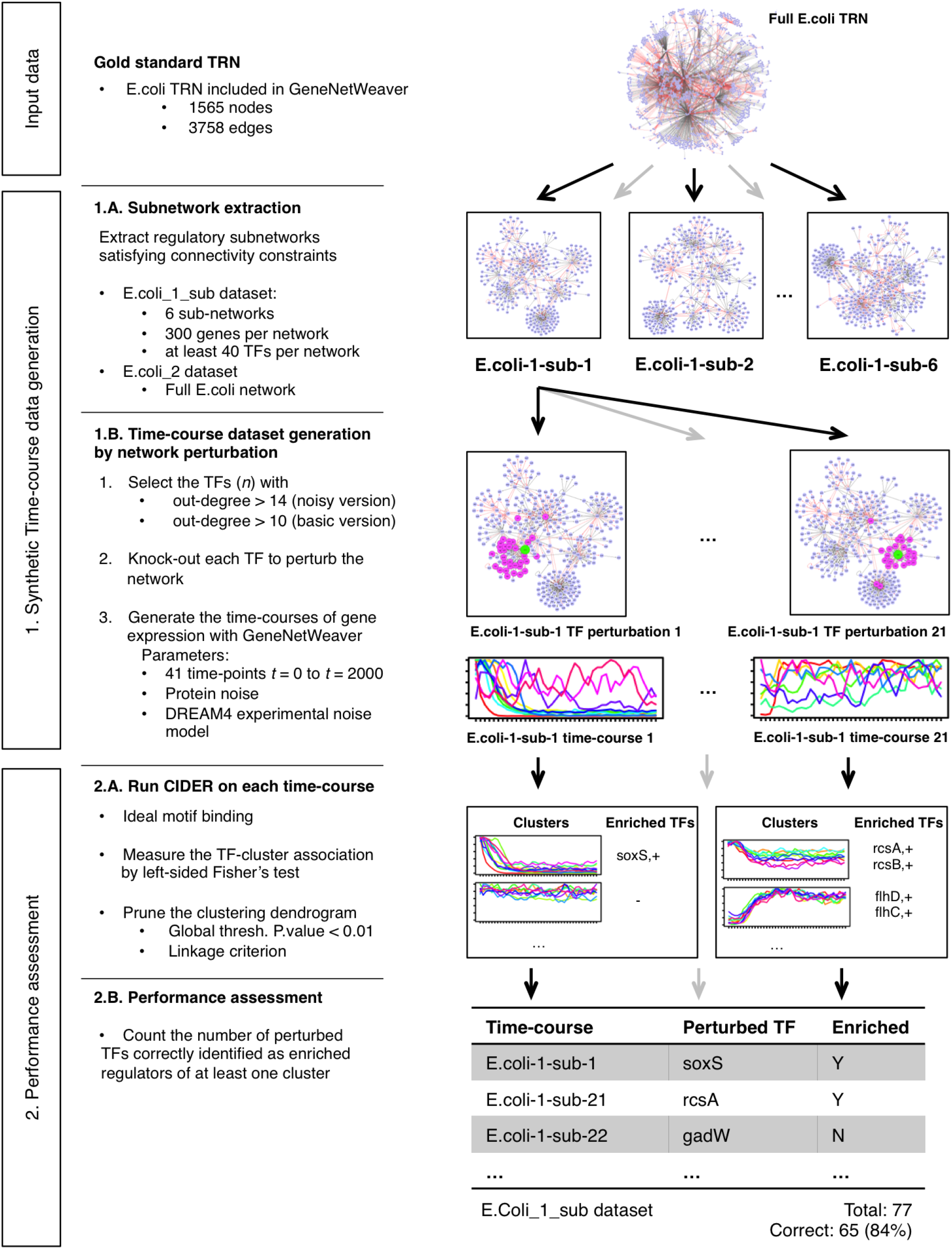
Description of the assessment strategy based on the two synthetic time-course gene expression datasets generated by GeneNetWeaver.

### 3.1 Description of the In-silico datasets and validation procedure

We generated two synthetic datasets by perturbing the *E.coli* transcriptional regulatory network and collecting the resulting gene expression time series. GeneNetWeaver (version 3.1b), a mature software to generate in-silico gene expression datasets given a known model of regulatory network [23], was used to induce perturbations in the known *E.coli* regulatory network included in the software (1565 genes, 3758 interactions), and simulate the resulting gene expression dynamics. In details, GeneNetWeaver was instructed to simulate a single knock-out event for each TF of the input network and register the resulting time series of gene expression for all the genes within the network (step 1.B in Fig. 2). All the resulting time-courses were automatically normalized by GeneNetWeaver between 0 and 1.

We observed that in some cases the expression variability of the knocked-out TF was not enough pronounced to induce a measurable effect on the regulatory network, leading to time-constant expression patterns for most of the genes. Hence, in order to build a dataset of timecourses with resonable signal variability, we only considered the time-courses generated by significant perturbations of the knocked-out TF. Precisely, a TF was considered significantly perturbed when its normalized expression varied of at least 0.25 across the entire time-course.

The two synthetic datasets differs in the size and complexity of the regulatory network considered:

- **Ecoli_1_sub dataset**: The first synthetic dataset collects time-courses derived from the perturbation of small regulatory networks. GeneNetWaver was used to extract six subnetworks of 300 genes (hereby referred to as ecoli_1_sub x) from the complete *E.coli* regualtory network (step 1.A in Fig. 2). The default subnetwork extraction parameters were used, and GeneNetWeaver was required to include at least 40 TFs in each subnetwork. In total we collected 157 time-courses associated to significant TF perturbations.
- **Ecoli_2_full dataset**: We built a second validation dataset considering the entire *E.coli* TRN (1565 nodes, 3758 edges). Gene expression time-courses were simulted by GeneNetWeaver following the same procedure used for the ecoli_1_sub dataset. In total we collected 139 time-courses associated to significant TF perturbations (out of the 178 TFs within the whole *E.coli* regulatory network).

The resulting gene expression dynamics was recorded in time-courses of 41 time-points, covering the time span between 0 (beginning of the knockout perturbation) and 2000 seconds. GeneNetWeaver produced two versions of each time-course: a “basic” version without experimental measurement noise added, and a “noisy” version with experimental noise added, following the noise model defined in the DREAM4 challenge [24]. We evaluated the performance of CIDER on both versions, thus providing an estimate of the sensibility of CIDER to noise. Note that both the basic and noisy time-courses were generated by using stochastic rather than ordinary differential equations, thus modeling the molecular noise effects on the network regulation.

For the multilevel motif enrichment analysis we set up an ideal motif binding scenario with exact information on binding activity, by exploiting the known structure of the *E.coli* regulatory network (step 2.B in Fig. 2). This was useful to assess the clustering procedure and the motif enrichment in an ideal scenario without motif enrichment bias. For each TF, only the differentially expressed target genes were used as background set in the enrichment analysis. The three enrichment pruning strategies were used altogether to yield the strictest results. The absolute enrichment P value was set to 0.01, and the standard deviation parameter *α* was fixed to 1.5.

### 3.2 Detectability threshold

Perturbing TFs elicits a stronger or weaker network response, measurable in terms of gene expression variability, depending on the extension and on the topology of the portion of regulatory network controlled by the TF. For instance, if a TF *x* regulates a small number of genes *Y*, the effect of perturbing *x* might elicit only a feeble variability in the regulatory network, not noticeable when looking at the expression dinamics. This phenomenon is even stronger when there are not TFs within the set *Y*. Another phenomenon that would limit the effects of the perturbation of *x* can potentially manifest when the downstream targets *Y* of *x* are also regulated by another TF *z*. If the regulatory strength of *z* on *Y* is stronger than *x*, then perturbing *x* might not be sufficient to induce a detectable effect in the network.

Beside these biological aspects, there are also technical limitations to the detectability of TF activity, as the one imposed by the use of the Fisher’s test in the motif enrichment analysis. Indeed, when the number of potential targets of a TF within the perturbed network is small, the estimation of the significativity by the Fisher’s test may not be accurate, resulting in not significant enrichment scores.

Hence, in order to provide indications on the applicability of CIDER, we studied the varibility of CIDER output respect to the degree of the perturbed TF. The TF degree is defined as the number of downstream targets that the TF possesses in the regulatory network used to generate the time-course. For instance, in the *E.coli* subnetworks, the degree of the TF is the number of genes regulated by the TF in the subnetwork, not on the entire *E.coli* network. Additionally, the number of differentially expression (DE) genes in the time-course was considered as covariate. As done for the TFs, a gene was considered as DE if its expression variability was *>* 0.25 across the entire timecourse.

The relationship between the degree of the perturbed TF, the number of DE genes, and the outcome of CIDER analysis for each time-course of the noisy ecoli_1_sub dataset is displayed in Fig. 3a. As expected, CIDER tends to return no enriched clusters for the time-courses associated to the perturbation of low-degree TFs. The number of DE genes in the time-course, instead, does not appear to influence the outcome. A similar pattern is observed when considering the basic version of the ecoli_1_sub dataset (Fig. S1a).

**Fig. 3:**
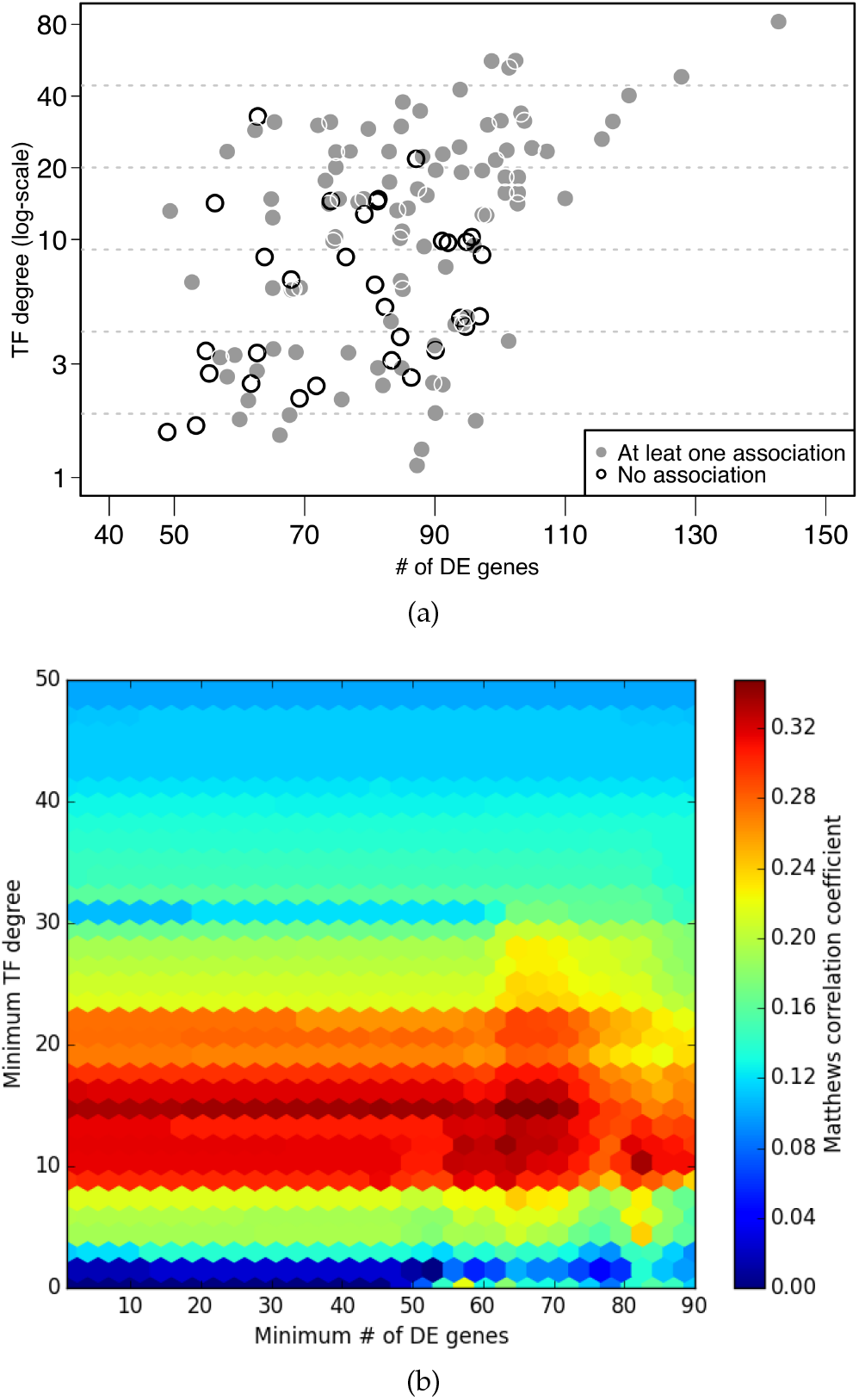
Variability of CIDER output respect to the degree of the perturbed TF and to the number of DE genes in the associated time-course, considering the 157 time-courses in the noisy version of the *E.coli_*1_sub dataset. (a) Solid gray dots represent time-courses for which CIDER reported at least one enriched TF. Empty black circles represent timecourses for which no enrichment was detected by CIDER. (b) MCC scores used in the grid search of the detectability threshold.

In order to empirically determine a “detectability threshold”, we performed a grid-search for the less stringent threshold, in terms of TF degree and number of DE genes, that best separate the time-courses for which CIDER is respectvely able and unable to detect any transcriptional wave. The optimization was performed by considering the set *N* of time-courses in the noisy version of the ecoli_1_sub dataset (|*N|* = 157). Given a pair *i* = (*x, y*) ∈ 𝒩^2^, let *T*_*i*_ *∈ N* be the subset of the time-courses that are associated to the knock-out of a TF with degree *> x* and have a number of DE genes *> y*. Let *D ∈N* be the subset of the time-courses for which CIDER identified at least one enriched cluster, regardless of its correctness. The set *D*_*i*_ = *D*∩*T*_*i*_ represents the time-courses satisfying the threshold *i* that have a signal detectable by CIDER. Conceptually, the optimal threshold should exclude most of the time-courses with no enrichment but retain most of those with at least one enrichment. We make use of the Matthew Correlation Coefficient (MCC) to evaluate the trade-off between including timecourses without enrichment and excluding those with sufficient signal. The optimal detectability threshold is the pair 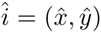 with highest MCC. The MCC for a generic threshold *i* is defined as:

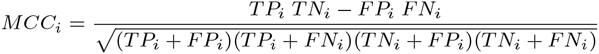

where

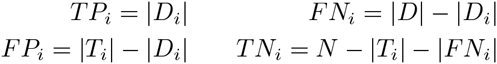

We found a ridge of thresholds with the similar maximum MCC at 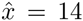, extending over a wide range of *y* (Fig. 3b). The presence of this ridge indicates that the number of DE genes in the time-course does not affect CIDER ability to detect transcriptional waves, while the TF degree is a significant factor. Consistently, we found a slightly less stringent optimal detectability threshold 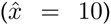 when considering the basic version of the ecoli_1_sub dataset (Fig. S1b).

The results of this section suggest that controlling at least 15 genes is the minimum requirement for eliciting a perturbation detectable by CIDER. We believe that this limit is mainly due to the Fisher’s test used in the motif enrichment analysis, hence improving the enrichment analysis might leverage this constraint. Note that this constraint is not problematic, as in the larger regulatory networks, such as those of human and mouse, the TFs generally regulate dozens of genes, extending safely beyond the detectability threshold.

### 3.3 Performance evaluation on the ecoli_1_sub synthetic dataset

For each time-course, the set of TFs found enriched in clusters of genes with specific expression patterns was compared to the list of knocked-down TFs. CIDER was considered to perform correctly when the perturbed TF or a TF directly regulated by the perturbed TF was included in the list of enriched TFs.

Only the 47 noisy and 77 basic time-courses associated to the significant perturbation of a TFs with degree greater than the detectability thresholds of 11 and 15 genes, respectively for the basic and noisy versions, were considered. For the basic ecoli_1_sub dataset, CIDER found the perturbed TF (or one of its direct downstream TFs) correctly enriched in 65 out of the 77 time-courses (84%, Table 1 and Fig. S2). The performance on the noisy version of the ecoli_1_sub dataset are comparable, with 38 correct inferences out of 47 considered time-courses (81%, Table 2 and Fig. 4).

**Fig. 4:**
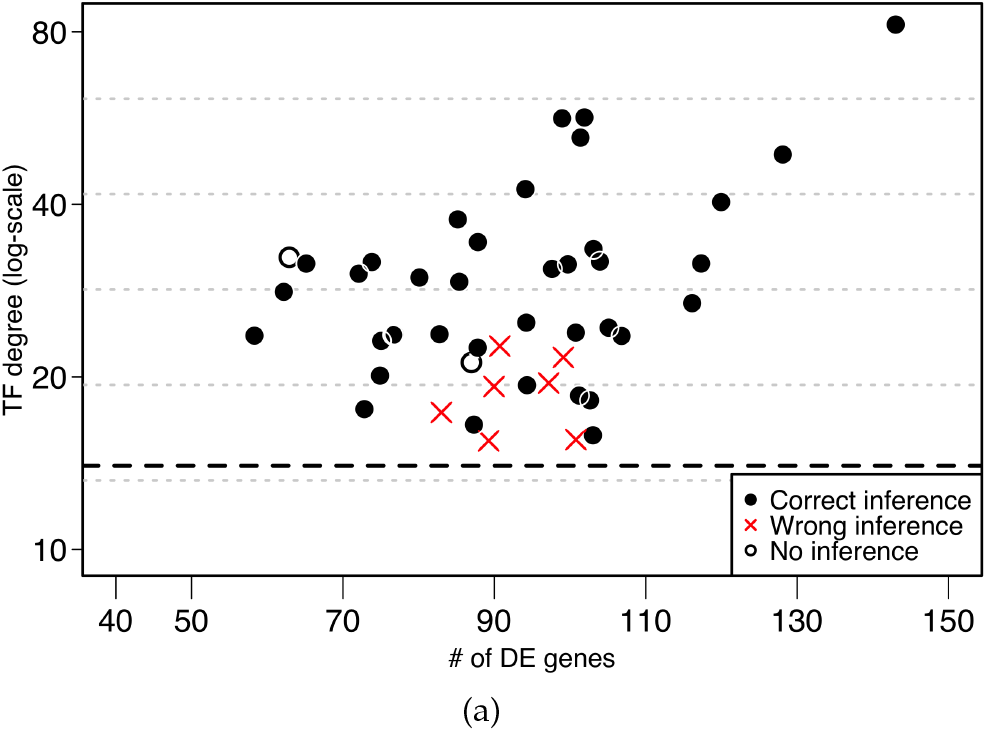
Performance of CIDER analysis on the noisy version of the *E.coli_*1_sub dataset. Solid black circles represent time-courses where CIDER correctly identified the perturbed TF (or a direct downstream TF) as regulator of a transcriptional wave. Red crosses represent time-courses where CIDER did not identify the perturbed TF (or a direct downstream TF) as regulator of a transcriptional wave. Empty black circles represent time-courses for which CIDER did not report any enrichment. The dashed line represent the detectability threshold (TF degree *≥* 15).

**Table 1:** Description of the noisy version of the Ecoli_1_sub dataset, and the performance of CIDER analysis.

| Network | Nodes | Edges | Number of Time-courses<sup>1</sup> | Correct inference # | % |
| --- | --- | --- | --- | --- | --- |
| Ecoli_1_sub_1 | 300 | 577 | 4 | 3 | 75 |
| Ecoli_1_sub_2 | 300 | 580 | 7 | 7 | 100 |
| Ecoli_1_sub_3 | 300 | 532 | 11 | 9 | 82 |
| Ecoli_1_sub_4 | 300 | 577 | 6 | 6 | 100 |
| Ecoli_1_sub_5 | 300 | 481 | 8 | 5 | 62 |
| Ecoli_1_sub_6 | 300 | 671 | 11 | 8 | 72 |
| Ecoli_1_sub (cumulative) |  |  | 47 | 38 | 81 |

<sup>1</sup> Number of time-courses associated to a significant perturbation (expression variability $> 0.25$ ) of a TF with degree $\geq 15$ .

### 3.4 Performance evaluation on the ecoli 2 synthetic dataset

Next, CIDER was validated on the Ecoli_2 dataset. The same validation approach was used, considering only the time-courses associated to 23 perturbed TFs with degree *≥*15 in both the basic and noisy versions of the dataset. CIDER correctly inferred the activity in 20 out of 23 timecourses (87%) in the noisy version, and of 27 out of 31 time-courses in the basic version (87%).

**Table 2:** Description of the basic version of the Ecoli_1_sub dataset, and the performance of CIDER analysis.

| Network | Nodes | Edges | Number of Time-courses<sup>1</sup> | Correct inference # | % |
| --- | --- | --- | --- | --- | --- |
| Ecoli_1_sub_1 | 300 | 577 | 10 | 10 | 100 |
| Ecoli_1_sub_2 | 300 | 580 | 13 | 13 | 100 |
| Ecoli_1_sub_3 | 300 | 532 | 15 | 12 | 80 |
| Ecoli_1_sub_4 | 300 | 577 | 13 | 9 | 69 |
| Ecoli_1_sub_5 | 300 | 481 | 11 | 9 | 81 |
| Ecoli_1_sub_6 | 300 | 671 | 15 | 12 | 80 |
| Ecoli_1_sub (cumulative) |  |  | 77 | 65 | 84 |

<sup>1</sup> Number of time-courses associated to a significant perturbation (expression variability $> 0.25$ ) of a TF with degree $\geq 11$ .

The scatter plots in Fig. 5 and Fig. S3 show the performance of CIDER on the noisy and basic versions of the Ecoli_2 datasets, respectively. Consistently with the detection limits derived form the noisy Ecoli_1_sub dataset, CIDER failed to predict the activity of most of the time-courses generated under the perturbation of TFs ≤ with degree 14.

Note that in this evaluation we considered as correct only the inferences that associate the knocked-out TF or one of its direct downstream TFs to a transcriptional wave. This does not mean that CIDER was unable to correctly identify any transcriptional wave in the timecourses marked as wrong in Fig. 5. A detailed example is provided in the case-study described in the next section.

### 3.5 Case study of CIDER analysis

This paragraph describes in details the CIDER analysis of the time-course associated to the knock-out of the *oxyR* TF in the Ecoli_1_sub_2 subnetwork. The noisy version of the time-course is considered. This time-course was not included in the validation described in the previous paragraphs, as *oxyR* was not significantly perturbed by GeneNetWeaver (its expression variability was less that the 0.25 threshold of DE). We selected this time-course to prove that even in this extreme case CIDER is able to detect some of the downstream effects of the *oxyR* perturbation and correctly identify the transcriptional waves induced in the regulatory network.

CIDER identified four transcriptional waves respectively regulated by *fur*, *metJ*, *flhC/D*, and *marA*, involving in total 32 of the 88 genes differentially expressed in the time-course (Table 3).

These inferred transcriptional waves were manually compared to the structure of the Ecoli_1_sub_2 network to determine their consistency with the effective propagation of the *oxyR* perturbation in the regulatory network. The portion of the Ecoli_1_sub_2 network interested by the perturbation is organized in Fig. 6, following the transcriptional cascade triggered by the knock-out.

**Fig. 5:**
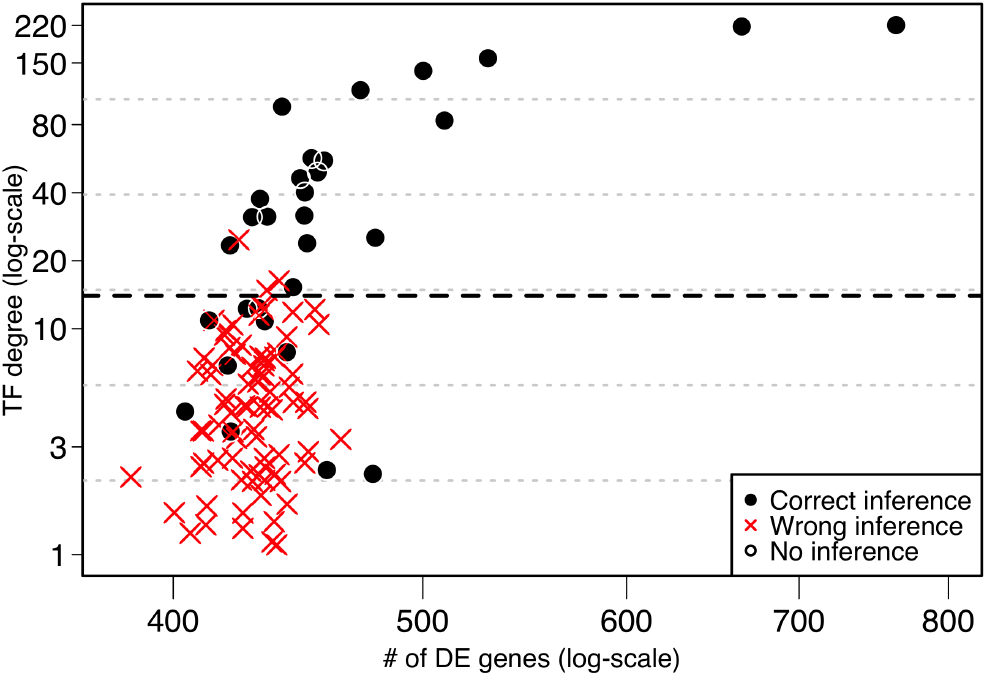
Performance of CIDER analysis on the noisy version of the *E.coli_*2 dataset. Solid black circles represent time-courses where CIDER correctly identified the perturbed TF (or a direct downstream TF) as regulator of a transcriptional wave. Red crosses represent time-courses where CIDER did not identify the perturbed TF (or a direct downstream TF) as regulator of a transcriptional wave. Empty black circles represent time-courses for which CIDER did not report any enrichment. The dashed line represent the detectability threshold (TF degree *≥* 15).

**Table 3:** Enrichment table produced by CIDER in the analysis of the noisy version of the time-course associated to the knock-out of the oxyR TF in the Ecoli_1_sub_2 network.

| Cl. Id | TF | Cluster size | Correct targets | Hit ratio | Fisher test P value |
| --- | --- | --- | --- | --- | --- |
| 1 | *fur*, - | 9 | 8 | 0.89 | $9.28 \cdot 10^{-6}$ |
| 2 | *metJ*, - | 7 | 4 | 0.57 | 0.0002 |
| 3 | *flhD*, + | 7 | 6 | 0.86 | $3.55 \cdot 10^{-5}$ |
| 3 | *flhC*, + | 7 | 6 | 0.86 | $3.55 \cdot 10^{-5}$ |
| 4 | *marA*, + | 9 | 5 | 0.56 | 0.0003 |

The *oxyR* knock-out induces the downregulation of the *fur* TF. As *fur* downregulates the *metJ*, *flhC* and *flhD* TFs, the perturbation initiated by *oxyR* is propagated downstream in the transcriptional network. Note that the strength of fur regulation on the downstream TFs is strongly variable, as confirmed by the significantly different expression patterns of *metJ* respect to *flhC/D*. CIDER was unable to detect any direct regulatory effect of *oxyR*, including the downregulation of *fur*, as the weak *oxyR* perturbation does not elicit a consistent effect on a relevent subset of its direct targets. Indeed, the set of *oxyR* direct targets encompasses only 15 genes that are regulated by different mechanisms involving multiple TFs, resulting in different expression patterns that are not clustered together by CIDER.

CIDER was able to identify three transcriptional waves that are induced downstream of *oxyR* by propagation of the perturbation across the transcriptional network. As the *fur* expression decreases, the expression of a first wave of genes negatively targeted by *fur* consistently increases (cluster 1, 9 genes). This wave includes the *metJ* TF; as the expression of *metJ* increases, it induces a transcriptional wave on its downstream target genes, correctly detected by CIDER (cluster 2, 7 genes). Simultaneously, the *flhC/D* TFs, not downregulated by *fur*, induce the expression of the third transcriptional wave, recognized by CIDER (cluster 3, 7 genes).

## 4 CONCLUSION

CIDER is a novel methodology for the inference of the transcriptional waves induced by the regulatory activity of the TFs, orthogonal to the current approaches of network inference. CIDER is not designed to infer the single regulatory interactions between TFs and target genes, but aims at identifying the role of the TFs in driving the waves of consistent transcription triggered in the cell by external stimuli.

In this work, we assessed CIDER on two realistic synthetic datasets based on the *E.coli* regulatory network.

A remarkable aspect of CIDER is that it requires only a time-course from a single biological replicate. Provided that the number and arrangement of time-points is sufficient to discern the expression patterns, CIDER opens the possibility of analyzing transcritpional regulation at single-sample resolution.

**Fig. 6:**
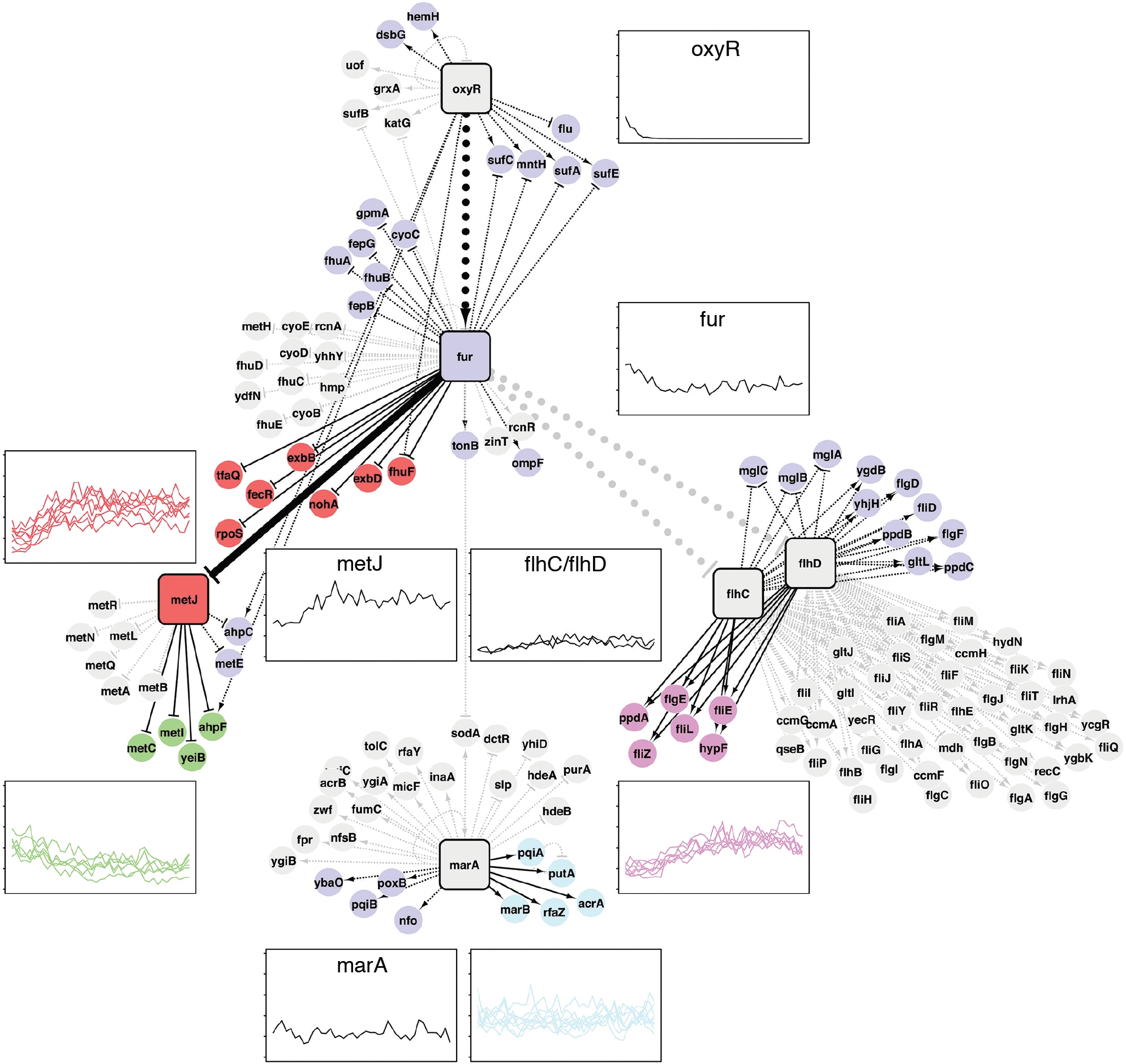
Transcriptional waves induced by the knock-out of oxyR in the Ecoli_1_sub_2 subnetwork, and correctly inferred by CIDER. Gray nodes represent not DE genes that are not considered in the analysis. Purple nodes represent DE genes that were not included in any enriched cluster. The other colored nodes (red, green, pink and light blue) represent genes included in clusters enriched for at least one TF. Squared nodes are TFs involved in the propagation of the perturbation or enriched in an expression cluster (oxyR, fur, metJ, flhC, flhD, marA). The black solid edges represent the transcriptional interactions correctly inferred by CIDER, the black dotted edges are regulatory interactions not captured by CIDER, and the gray dotted edges are regulatory interactions invoving genes not DE genes excluded from the analysis. Arrow-ending and flat-ending edges represent positive and negative regulation, respectively.

